# Inflammation in the COVID-19 airway is due to inhibition of CFTR signaling by the SARS-CoV-2 Spike protein

**DOI:** 10.1101/2022.01.18.476803

**Authors:** Hung Caohuy, Ofer Eidelman, Tinghua Chen, Qingfeng Yang, Bette S. Pollard, Nathan I. Walton, Harvey B. Pollard

## Abstract

**Background:** SARS-CoV-2-contributes to sickness and death in COVID-19 patients partly by inducing a hyper-proinflammatory immune response in the host airway. This hyper-proinflammatory state involves activation of signaling by NFκB and ENaC, and expression of high levels of cytokines and chemokines. Post-infection inflammation may contribute to “Long COVID”, and there are long term consequences for acute severe COVID-19, which double or triple the chances of dying from any cause within a year. Enhanced signaling by NFκB and ENaC also marks the airway of patients suffering from cystic fibrosis, a lethal proinflammatory genetic disease due to inactivating mutations in the CFTR gene. We therefore hypothesized that inflammation in the COVID-19 airway might be due to inhibition of CFTR signaling by SARS-CoV-2 Spike protein.

**Methods:** This hypothesis was tested using the hTERT-transformed BCi-NS1.1 basal stem cell, previously derived from small airway epithelia, which were differentiated into a model of small airway epithelia on an air-liquid-interface (ALI). CyclicAMP-activated CFTR chloride channel activity was measured using an Ussing Chamber. Cell surface-CFTR was labeled with the impermeant biotin method.

**Results:** Exposure of differentiated airway epithelia to SARS-CoV-2 Spike protein resulted in loss of CFTR protein expression. As hypothesized, TNFα/NFκB signaling was activated, based on increased protein expression of TNFR1, the TNFα receptor; TRADD, the first intracellular adaptor for the TNFα/TNFR1 complex; phosphorylated IκBα, and the chemokine IL8. ENaC activity was also activated, based on specific changes in molecular weights for α and γ ENaC. Exposure of the epithelia to viral Spike protein suppressed cAMP-activated CFTR chloride channel activity. However, 30 nM concentrations of cardiac glycoside drugs ouabain, digitoxin and digoxin, prevented loss of channel activity. ACE2 and CFTR were found to co-immunoprecipitate (co-IP) in both basal cells and epithelia, suggesting that the mechanism for Spike-dependent CFTR loss might involve ACE2 as a bridge between Spike and CFTR. In addition, Spike exposure to the epithelia resulted in failure of endosomal recycling to return CFTR to the plasma membrane, suggesting that failure of CFTR recovery from endosomal recycling might be a mechanism for spike-dependent loss of CFTR.

**Conclusion:** Based on experiments with this model of small airway epithelia, we predict that inflammation in the COVID-19 airway may be mediated by inhibition of CFTR signaling by SARS-CoV-2 Spike protein, thus inducing a CFTR-null, cystic fibrosis-like clinical phenotype.

## Introduction

SARS-CoV-2-contributes to sickness and death in COVID-19 patients partly by inducing a hyper-proinflammatory immune response in the host airway ^1, 2^. Consistently, COVID-19 patients who have been admitted to the Intensive Care Unit (ICU) have the most severe forms of inflammatory disease ^3^. This hyper-proinflammatory state, also termed cytokine storm ^4^, or cytokine release syndrome ^5^, involves activation of signaling by NFκB ^6, 7^ and by ENaC, the Epithelial Sodium channel ^8, 9^. Sustained, post infection inflammation may also contribute to Post-Acute Sequelae of COVID-19 (PASC), also termed “long COVID” ^10–12^, and to acute “Severe COVID-19”, which doubles or triples the chances of dying from any cause within a 6-12 month period ^13, 14^. However beyond the specific involvement of NFκB and ENaC in COVID-19, the actual mechanisms remain poorly understood. NFκB, the master regulator of inflammation, contributes to cytokine storm by driving the synthesis and secretion of cytokines and chemokines ^15^. In response to NFκB activation the COVID-19 airway responds to IL-8 and other cytokines by attracting high levels of neutrophils ^16^, and changes in other types of immune cells ^16, 17^. Activation of ENaC in the airway also contributes to inflammation, as well as to airway surface dehydration and impaired mucociliary clearance ^9, 18, 19^. SARS-CoV-2 infection is initiated by the viral Spike protein binding to the cellular receptor ACE2. However it remains to be better understood how inflammation follows from this interaction. We have recently reported that nanomolar concentrations of cardiac glycosides such as ouabain, digitoxin and digoxin competitively inhibit binding of the SARS-CoV-2 Spike protein to the ACE2 receptor ^20^. Consequently they also block both penetration by Spike-pseudotyped virus and infectivity by native SARS-CoV-2 in human lung cells ^20^. Thus these candidate anti-viral drugs could additionally serve as tools to help elucidate the virus-specific pro-inflammatory mechanisms.

Coincident activation of both NFκB and ENaC in the airway is also a characteristic of cystic fibrosis (CF), a rare, lethal, autosomal recessive genetic disease. In this case the proinflammatory activation mechanisms for both NFκB and ENaC depend on the presence of inactivating mutations in the Cystic Fibrosis Transmembrane Conductance Regulator protein CFTR ^9, 21–24^. Under resting conditions, CFTR sends constitutively synthesized TRADD, the first intracellular adaptor of the TNFα/TNFR1 complex, to the proteosome ^25^ (see also **Supplemental Figure S1**). Without TRADD activation, the TNFα/NFκB signaling pathway is silent and NFκB p65 remains inactively complexed with IκBα in the cytosol. However, in the absence of CFTR, TRADD survives in the cytosol to signal to the IKKα,b,g complex to phosphorylate IκBα and send p- IκBα to the proteosome. NFκB,p65 is thus freed to enter the nucleus and to drive proinflammatory cytokine and chemokine expression. Similarly, although by a less well understood mechanism, CFTR constitutively suppresses ENAC activity in the lung ^26, 27^. Because activated ENaC removes sodium from the airway, the purpose of this suppression mechanism is understood to ensure adequate hydration of the airway. Coincidentally, in response to active Cl- conductance by the cAMP-activated CFTR chloride channel, Na+ is passively conducted into the airway through inactive ENaC channels. Water is then passively moved into the airway in response to the resulting osmotic gradient. However, in the absence of CFTR-mediated chloride movement into the airway, ENaC is proteolytically activated by TMPRSS2 and FURIN. The now activated ENaC channel uncontrollably dehydrates the airway by removing sodium from the airway and expelling it through the basolateral part of the cell ^27–29^. The airway responds to this dehydration damage with NFκB-dependent inflammation, including activation of the NLRP inflammasome and cytokines such as IL-18, IL-1β and caspase 1 ^30^. ENaC in the lung is simultaneously activated by Angiotensin II which is synthesized in the lung by ACE in the pulmonary vasculature, and inactivated by epithelial ACE2 ^31, 32^ **(see Supplemental Figure S1).** Conceptually, these interactions are part of a massive CFTR interaction network, which was first understood from the viewpoint of interlocking PDZ-based molecular switches ^33^. Later the CFTR network was identified with multiple molecular chaperones and other functional proteins ^34, 35^. More recently the CFTR network has been identified in terms of a complex set of multidimensional systems biology-based interactive maps for wildtype and mutant CFTR ^24^. Thus CFTR is a key regulator of airway hydration and control of proinflammatory TNFα/NFκB and ENaC signaling.

Based on this close proinflammatory relationship in the airway between COVID-19 and Cystic Fibrosis (CF) symptoms, and by support from preliminary data, we have hypothesized that inflammation in the COVID-19 airway might be due to inhibition of CFTR signaling by the SARS-CoV-2 Spike protein. However, there has been conflicting data regarding the role of the viral Spike protein in driving proinflammatory signaling in cultured cells ^36, 37^. Therefore as a more physiological platform to test this hypothesis we used the hTERT-transformed BCi-NS1.1 basal stem cell, isolated from the small airways epithelium, which we studied in differentiated form as a model airway epithelium at the air-liquid-interface (ALI) ^38–40^. Attractive features of this model epithelium include (i) histologic and molecular fidelity to normal small airway epithelial cell physiology ^38, 39^; (ii) a normal male karyotype, 46,X/Y ^38–40^; (iii) cell-specific expression of COVID- 19 related genes including ACE2, ADAM 10 and ADAM 17, TMPRSS2, FURIN, and CTSL ^38^; and (iv) expression of CFTR protein by both basal cells and the differentiated epithelia (our preliminary data). To test the hypothesis we have asked whether exposure of the differentiated epithelium to SARS-CoV-2 Spike protein (1) activates critical parts of the TNFα/NFκB and ENaC signaling pathways; (2) induces loss of CFTR protein and cAMP-activated CFTR chloride channel activity; (3) rescues loss of CFTR channels by the cardiac glycoside drugs, ouabain, digitoxin and digoxin; and (4) suppresses CFTR levels by inhibiting endosomal recycling of CFTR. We also tested whether (5) epithelial ACE2 binds to CFTR. We found positive answers to all five tests, and concluded that the origins of the proinflammatory COVID-19 airway may be traced in part to the ability of the Spike protein to induce an approximately CFTR-null phenotype in the virus infected lung. *To our knowledge this is the first time COVID-19 airway inflammation has been experimentally traced to a contribution from SARS-CoV-2 Spike-dependent inhibition of CFTR signaling*.

## Results

### SARS-CoV-2 Spike protein activates TNFα/NFκB signaling in airway epithelium

As described above, CFTR tonically blocks activation of the TNFα/NFκB signaling pathway by destructively interacting with TRADD, the <u>T</u>NF <u>A</u>ssociated <u>D</u>eath <u>D</u>omain protein ^25^. Therefore, to determine whether the inhibitory CFTR/TRADD interaction is reduced in Spike-treated epithelia, we tested whether expression of TRADD was increased when epithelia were exposed to the Spike protein from the original strain of SARS-CoV-2 isolated in 2019 in Wuhan, China (Original-[S1S2] Spike). In parallel we also tested whether other proteins in the TNFα/NFκB signaling pathway, such as TNFR1, p-IκBα and the chemokine IL-8, were also increased. **Figure 1a** shows that as Spike protein increases, so do the protein expressions for TRADD, TNFR1 and p-IκBα. **Figure 1b** also shows that IL-8, as measured in the liquid sub- phase, increases in proportion to increases in Spike protein concentrations applied to the epithelia. Thus the entire proinflammatory TNFα/NFκB signaling pathway, including TRADD, is activated by the Spike protein. *Inasmuch as TRADD is increased by Spike exposure, it is therefore possible that loss of CFTR function could be involved in the Spike-dependent activation of proinflammatory TNFα/NFκB signaling in airway epithelia*.

**Figure 1.**
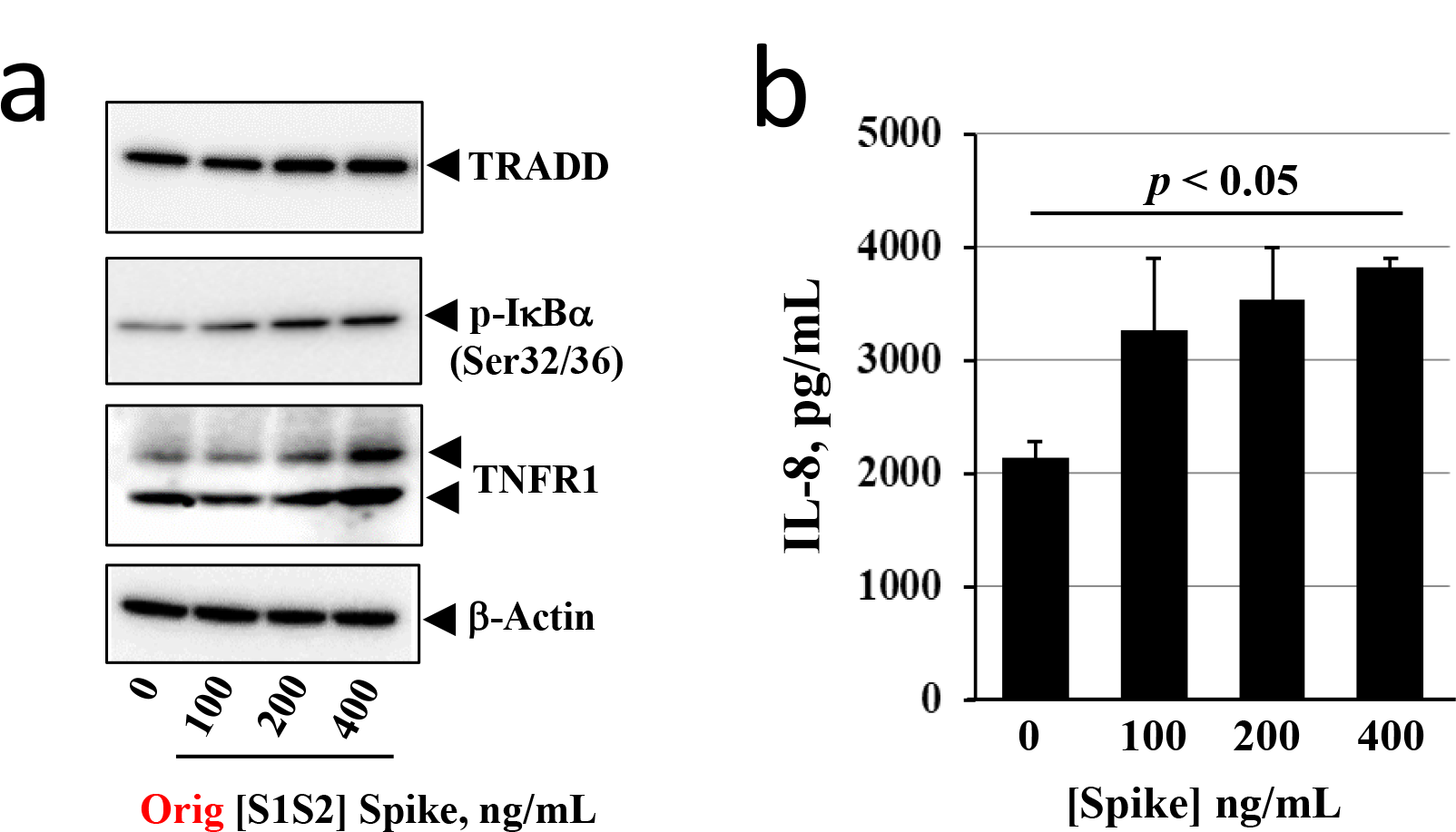
Effects of SARS-CoV-2 [S1S2] Spike on pro-Inflammatory TNFα/NFκB signaling proteins and IL-8 expression in differentiated BCi.NS1.1 (d-BCi) epithelia. The apical surface of the differentiated epithelium was treated with increasing concentrations of Original-[S1S2] Spike protein for 4 hours in submerged conditions, washed and then incubated for an additional 20 hours in ALI conditions. (a). TRADD, TNFR1 and p-IκBα increase in a Spike concentration-dependent manner. (b). IL-8 expression significantly increases as a function of increasing concentration of Spike protein. (N=3 independent experiments; *p* < 0.05 taken as significantly different from control.

### Spike protein reduces CFTR chloride channel activity in differentiated airway epithelia

To test whether Spike proteins affect CFTR channel function, we analyzed Spike-treated epithelia with the Ussing Chamber method. **Figure 2a** shows that under control conditions, cAMP-activated CFTR chloride channels can be detected, and can be specifically blocked by the CFTR channel inhibitor CFTRinh-172. However, when Original-[S1S2] Spike protein was pre-incubated for four hours at the apical surface of the differentiated epithelial culture, CFTR channel activity was significantly and dose-dependently reduced when assayed 20 hours later. The inhibition constant (Ki) is 446 ng/ml for the Original-[S1S2] Spike protein (see **Table 1**).

**Figure 2.**
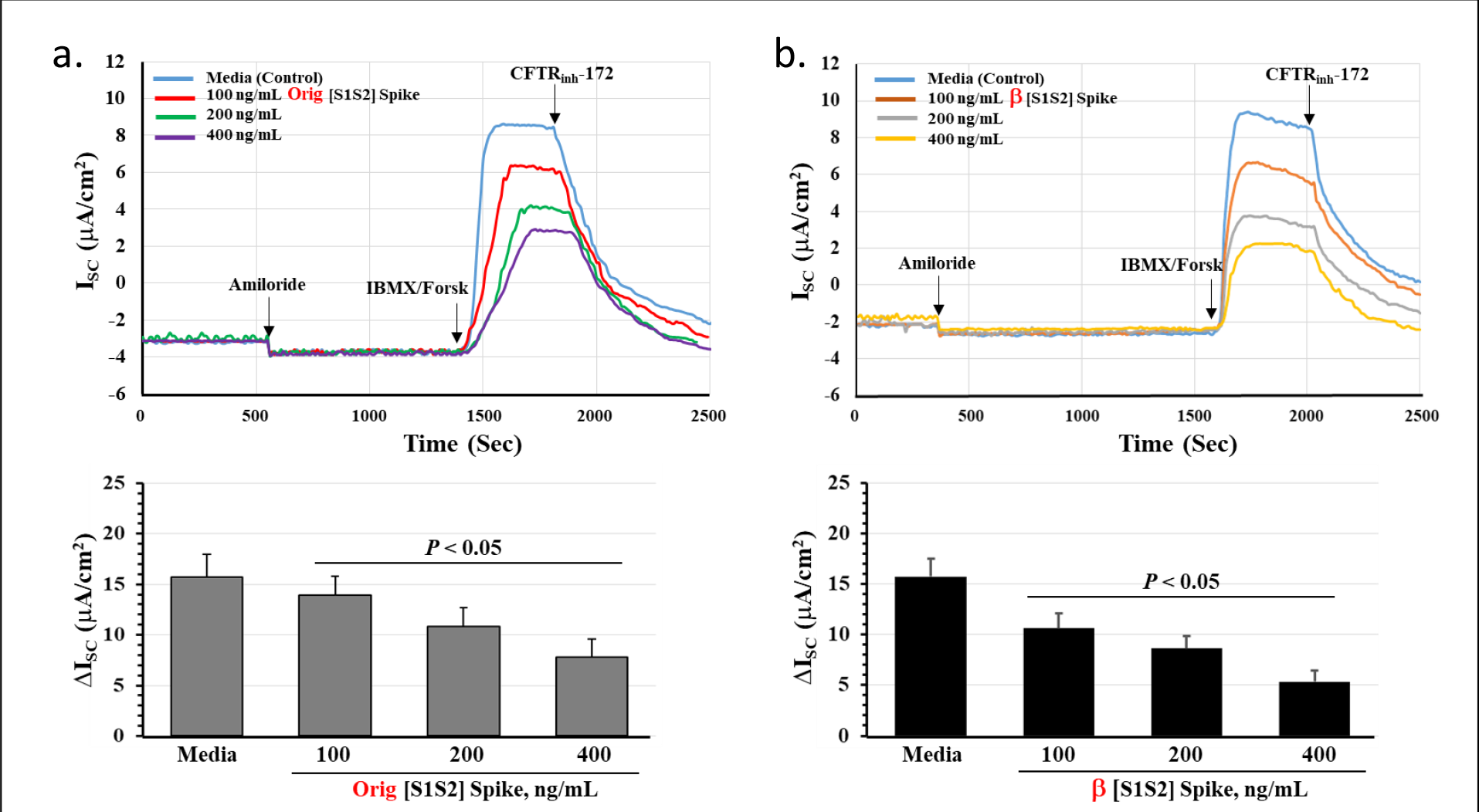
Original and. β **Spike Protein dose-dependently reduce cAMP-activated CFTR chloride channel activity. (a)** Different concentrations of Original-[S1S2] Spike protein from the original SARS-CoV-2 strain were incubated with epithelia for 4 hours in submerged conditions, washed and then incubated in medium for an additional 20 hours in ALI conditions. Chloride channels were analyzed in Ussing Chambers **(b)** Different concentrations of β-1.315 [S1S2] Spike protein were incubated and analyzed as in part **(a**.) (N=4 ± SE; *, P<0.05 compared to media control). Amiloride blocks Na+ channels. IBMX/Forsk activates cAMP synthesis. CFTRinh172 blocks CFTR Cl- channels.

**Table 1.** Inhibition constants for α and β Spike effects on CFTR channels and CFTR protein, and for binding constants for ACE2.

| **Spike Variant** | **K<sub>i</sub>, CFTR channel (R<sup>2</sup>)**<br>**ng Spike/ml** | **K<sub>i</sub>, CFTR protein(R<sup>2</sup>)**<br>**ng Spike/ml** | **K<sub>D</sub>, ACE2 (R<sup>2</sup>)**<br>**ng ACE2 /ml** |
| --- | --- | --- | --- |
| **Original-[S1S2]** | 446 (0.9898) | 399 (0.9842) | 318 (0.9960) |
| **β-1.315 [S1S2]** | 210 (0.9956) | 193 (0.9590) | 147 (0.9962) |
| **Ratio, %:**<br>**Original/β-1.315** | **(47%)** | **(48%)** | **(45%)** |

To test for the specificity of the Spike effect, we replaced the Original-[S1S2] Spike protein in the assay with the higher affinity β-1.315 [S1S2] Spike protein. As shown in **Figure 2b**, the β-1.315 [S1S2] Spike also reduced CFTR channel activity. However, the inhibition constant is 210 ng/ml for the β-1.315 [S1S2] Spike protein, or 47% lower than for the Original-[S1S2] Spike protein (see **Table 1**). These [S1S2] Spike proteins also differ in their affinity, or KD, for ACE2. **Table 1 and Supplemental Figure S2** show that the KD for ACE2 binding to the Original-[S1S2] Spike is 318 ng/ml ACE2, as compared with 147 ng/ml ACE2 for binding to the β-1.315 [S1S2] Spike. This 47% reduction for Spike binding parallels the approximately 45% lower KD value for ACE2 binding to the β-1.315 [S1S2] Spike protein. These data thus indicate that the relative KD values for Original- and β−1.315 Spike variants for inhibiting Channel activity and their respective affinities for ACE2 are very similar. *A possible interpretation is that the Spike proteins first bind to ACE2, and that the Spike:ACE2 complex may affect consequent CFTR function*.

### Cardiac glycoside drugs block Spike-dependent loss of CFTR channel activity

Cardiac glycosides such as ouabain, digitoxin and digoxin have been shown to be potent competitive inhibitors of ACE2 binding to the SARS-CoV-2 Spike protein ^20^. The proportional changes in CFTR channel inhibition constants for Original-[S1S1] Spike and β-1.315 [S1S2] Spike proteins, and respective binding constants to ACE2, suggest that CFTR channel reduction might be initiated by Spike proteins binding to ACE2 (see **Table 1)**. If this suggestion were correct then we could predict that the cardiac glycosides should block Spike-induced reduction in CFTR channel activity. As predicted, **Figure 3a** shows that 30 nM concentrations of each cardiac glycoside protects CFTR channels from reduction when epithelia are incubated with Original- [S1S2] Spike proteins. Further, **Figure 3b** reveal an apparent activity series of [ouabain, digitoxin > digoxin], which is similar to that reported for relative potencies as blockers of Spike:ACE2 binding ^20^. Based on both kinetic data and the use of cardiac glycosides as a tool to identify Spike:ACE2 binding, *it is increasingly likely that a direct Spike interaction with ACE2 is initially responsible for loss of CFTR channel activity*.

**Figure 3.**
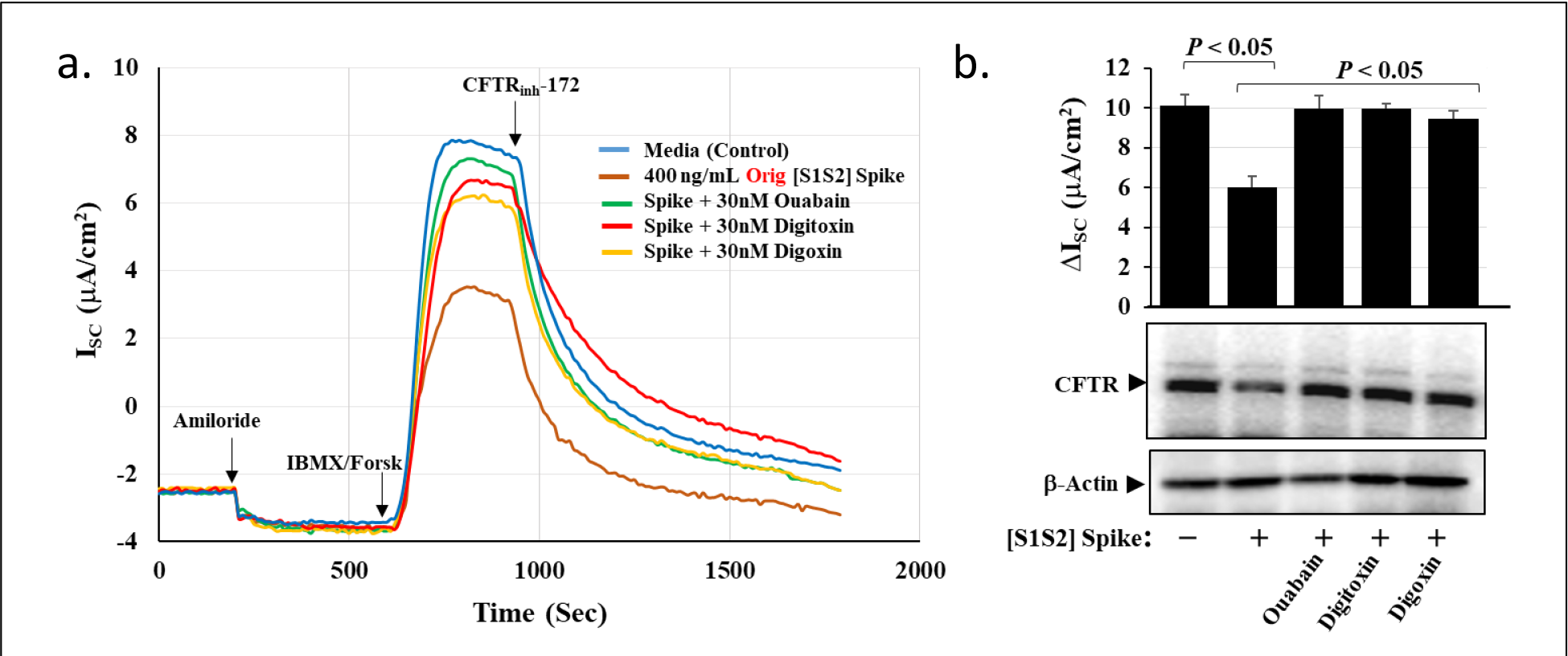
Cardiac glycosides block the inhibitory effect of SARS-CoV-2 [S1S2] Spike on cAMP-activated CFTR chloride channels in differentiated BCi.NS1.1 (d-BCi) epithelia. (a). Different concentrations of Original-[S1S2] Spike protein were incubated with epithelia for 4 hours in submerged conditions in the presence of 30 nM Ouabain, digitoxin or digoxin, washed and then incubated in in the ALI condition for an additional 20 hours. Chloride channels were analyzed in Ussing Chambers. **(b).** Data show that all three cardiac glycoside drugs significantly rescue the spike-dependent loss of channel activity. In addition, the second CFTR gel from the left shows that in the Spike/media-only control lane, CFTR protein appears to be reduced. (N=3 ± SE; *, P<0.05 compared to media control). Amiloride blocks Na+ channels. IBMX/Forsk activates cAMP synthesis. CFTRinh172 blocks CFTR Cl- channels).

Spike protein reduces CFTR protein expression in differentiated airway epithelia.

Scrutiny of the CFTR gels in **Figure 3b** reveals that exposure of the differentiated epithelia to Spike proteins appears to *reduce* the total CFTR in the system (see **Figure 3** legend for details). To test whether Spike-dependent loss of CFTR chloride channel activity might be due to loss of CFTR protein we examined the consequences of titrating both Original-[S1S2] Spike and β-1.315 [S1S2] Spike proteins on levels of total epithelial CFTR. **Figure 4a** and **b** show that both spike strains reduce total CFTR. However, the β-1.315 [S1S2] Spike protein appears to be the more potent. By comparing these data with those in **Figures 2a** and **b**, we noted that the patterns of reduction as a function of titrations of both Original and β spike proteins were remarkably similar to those for suppression of the cAMP-activated CFTR chloride channel. The Ki values for Original- [S1S2] Spike and β-1.315 [S1S2] Spike proteins, respectively, are 399 ng/ml and 193 ng/ml. The Ki value for β-1.315 [S1S2] Spike protein is thus reduced relative to the Original-[S1S2] Spike protein by 48% (see **Table 1**). The calculations of the kinetic values for CFTR channel and protein are shown in **Figure 5** and are summarized in **Table 1**. *The loss of CFTR channels may therefore be due to primary loss of CFTR protein*.

**Figure 4.**
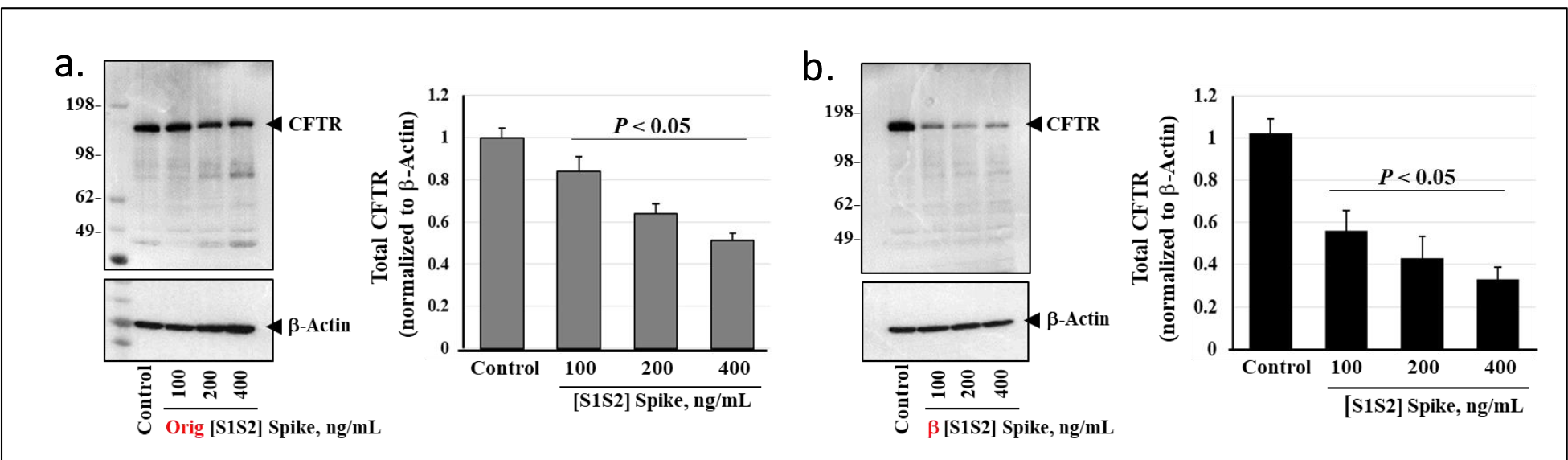
Original-[S1S2] Spike and. β**-1.315 [S1S2] Spike proteins inhibit apical and total CFTR protein expression in differentiated BCi.NS1.1 (d-BCi) epithelia. (a)** Different concentrations of Original-[S1S2] Spike protein were incubated with epithelia for 4 hours in submerged conditions, washed and then incubated in medium for an additional 20 hours in ALI conditions. Total CFTR protein in epithelia were measured by Western blot. **(b)** Different concentrations of β-1.315 [S1S2] Spike protein were incubated and analyzed as in part **(a**.) (N=4 ± SE; *, *p* < 0.05 compared to media control).

**Figure 5.**
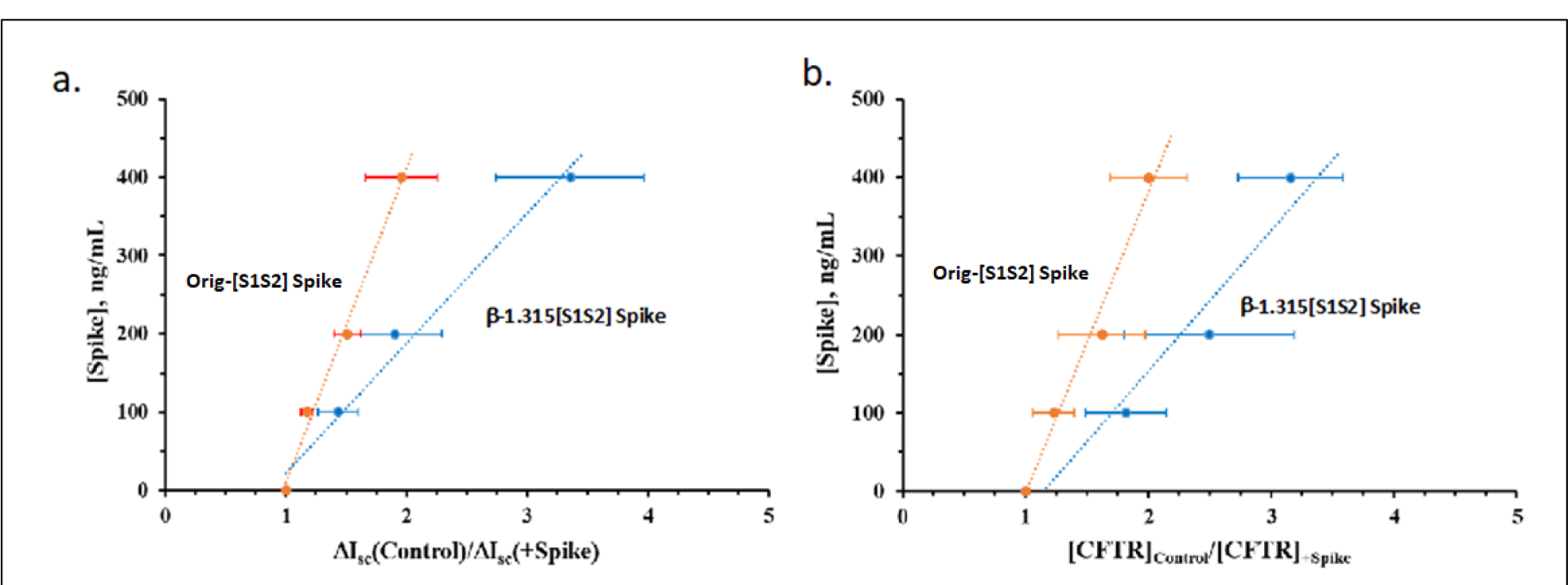
Ki values for cAMP-dependent CFTR chloride channel activity and for CFTR protein levels as a consequence of exposure of differentiated cultures to Original or. β **Spike proteins. (a)** CFTR Channel conductance as a function of Original and β Spike concentrations. Ki for Original-[S1S2] Spike is 448 ng/ml (R2 = 0.9898). Ki for β−Spike is 210 ng/ml (R2 =0.9956). (**b).** CFTR protein level as a function of Original and β Spike concentrations. Ki for Original−Spike is 399 ng/ml (R2 =0.9842) . Ki for β -Spike is 193 ng/ml (R2 = 0.9590). Graphs are derived from the Eadie-Hoffstee equation. Ki values are the slopes of the straight lines. Points are averages from N=4 ± SE independent experiments.

### CFTR co-immunoprecipitates with ACE2 in basal (b-BCi) cells and differentiated (d-BCi) epithelia

A critical question is how Spike proteins affect CFTR survival in epithelia when they must first bind to ACE2. One possibility is that ACE2 might serve as a physical bridge between the Spike protein and CFTR. To test this possibility, we asked whether CFTR and ACE2 could co-immunoprecipitate with each other. **Figure 6a** shows that both basal cells and differentiated epithelia contain ACE2 and CFTR. ACE2 has been previously reported to be in these basal cells, and it has been previously noted in native airways that aside from infrequent ionocytes, basal cells are second only to secretory cells in CFTR content ^41^. **Figure 6b** shows that if ACE2 were first immunoprecipitated from either basal cells or epithelia, both immunoprecipitates would contain CFTR. *Thus the possibility of ACE2 being a physical bridge, direct or indirect, between Spike protein and CFTR cannot be excluded*.

**Figure 6.**
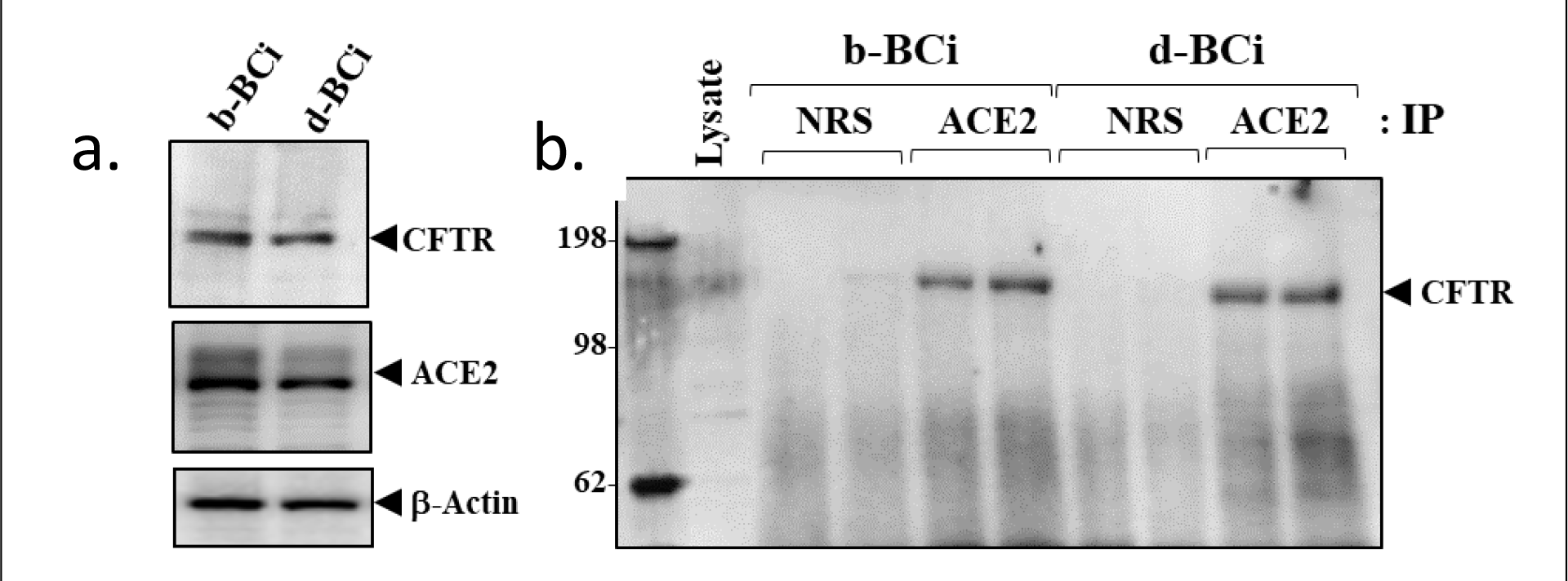
ACE2 binds CFTR in basal (b-BCi) cells and differentiated (d-BCi) epithelia. (a) Total CFTR and ACE2 levels in basal (b-BCi) and differentiated (d-BCi) epithelia. (**b**) CFTR bound to ACE2 in basal (b-BCi) and differentiated (d-BCi) epithelia.

### Spike protein inhibits CFTR recovery from endosomal recycling

If the binding of spike protein to ACE2 were a necessary precursor to the Spike-induced loss of CFTR from the plasma membrane, the next critical question would be how CFTR was physically removed and destroyed. Under normal conditions, CFTR on the plasma membrane is subject to constitutive endosomal recycling in which misfolded or otherwise damaged CFTR is diverted for destruction to the lysosome (see **Supplemental Figure S3**). By this mechanism approximately 33% of damaged or misfolded CFTR is lost to the lysosome in 10 minutes, while the remaining 67% is returned to the apical plasma membrane ^42^. To test whether spike-induced loss of apical CFTR was due to failure of recovery from endosomal recycling, we used an impermeant biotin labelling method to ask how much CFTR from the apical surface is returned to the apical surface after exposure to progressively increasing concentrations of Original-[S1S2] Spike protein. **Figure 7** shows that as the concentration of Original-[S1S2] Spike increases there is a steady reduction in recovery of recycled CFTR. This mechanism for CFTR loss due to exposure to Spike protein is thus consistent with failure of endosomal recycling to return initially apical CFTR to the apical plasma membrane. *It is therefore possible that when Spike protein binds to the ACE2:CFTR complex, the bound CFTR may become marked as “damaged ”* and thus *directed to the lysosome*.

**Figure 7.**
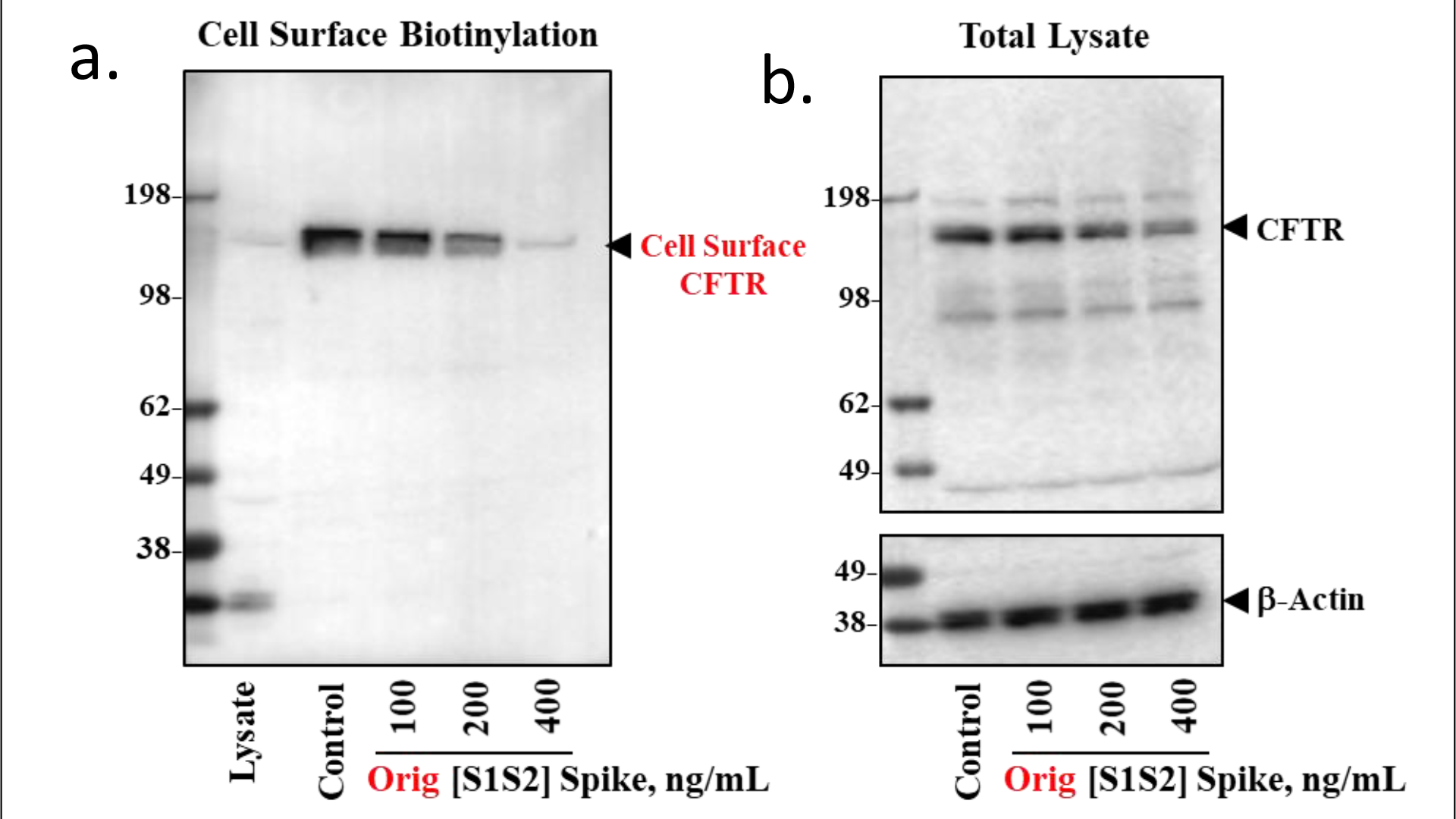
Effects of SARS-CoV-2 [S1S2] Spike on the endosomal recycling of cell surface CFTR protein expression in differentiated BCi-NS1.1 (d-BCi) Cells. (a) Endosomal recycling of cell surface CFTR by cell surface biotinylation as a function of exposure to increasing concentrations of Original-[S1S2] Spike protein. Epithelia were exposed to different concentrations of Spike protein for four hours; washed and incubated for a further 20 hours; then washed with ice-cold Ca2+/ Mg2+ PBS buffer 3 times; then incubated with 2 mg/ml impermeant biotinylation reagent sulfo-NHS-S-S biotin for 24 hours at 4oC before extraction with Strepavidin beads and assay by Western blot for CFTR. **(b)** Total lysates from the experiment in **Part a.** Note that total CFTR protein is also in decline as a function of concentration increases in original-[S1S2] Spike. This is consistent with the concept in **Supplemental Figure S3**, showing most CFTR flows to the plasma membrane through the TGN, and is endosomally recycled, either to the lysosome, the TGN or back to the plasma membrane. .

### Spike protein induces proteolytic activation of ENaC

Activation of ENaC occurs when the constitutive inhibitory activity of CFTR is lost, and sodium conductance is activated by proteolytic cleavage of the α and γ chains of the heterotrimeric ENaC channel ^9, 18, 19^. TMPRSS2, FURIN and possibly other serine proteases are responsible ^43, 44^. To test whether Spike protein treatment could activate ENaC, we asked whether proteolytic fragments of activated ENaC subunits could be detected following treatment of differentiated epithelial cells with Spike Protein. **Figure 8** shows that over the 20 hour period following the 4 hour exposure to Spike proteins, specific proteolytic fragments are detected for both α ENaC and γ ENaC. Furthermore, additional γ ENaC appears to have been synthesized and activated. *We conclude that Spike-dependent loss of CFTR is accompanied by proteolytic activation of both α ENaC and γ ENaC*.

**Figure 8.**
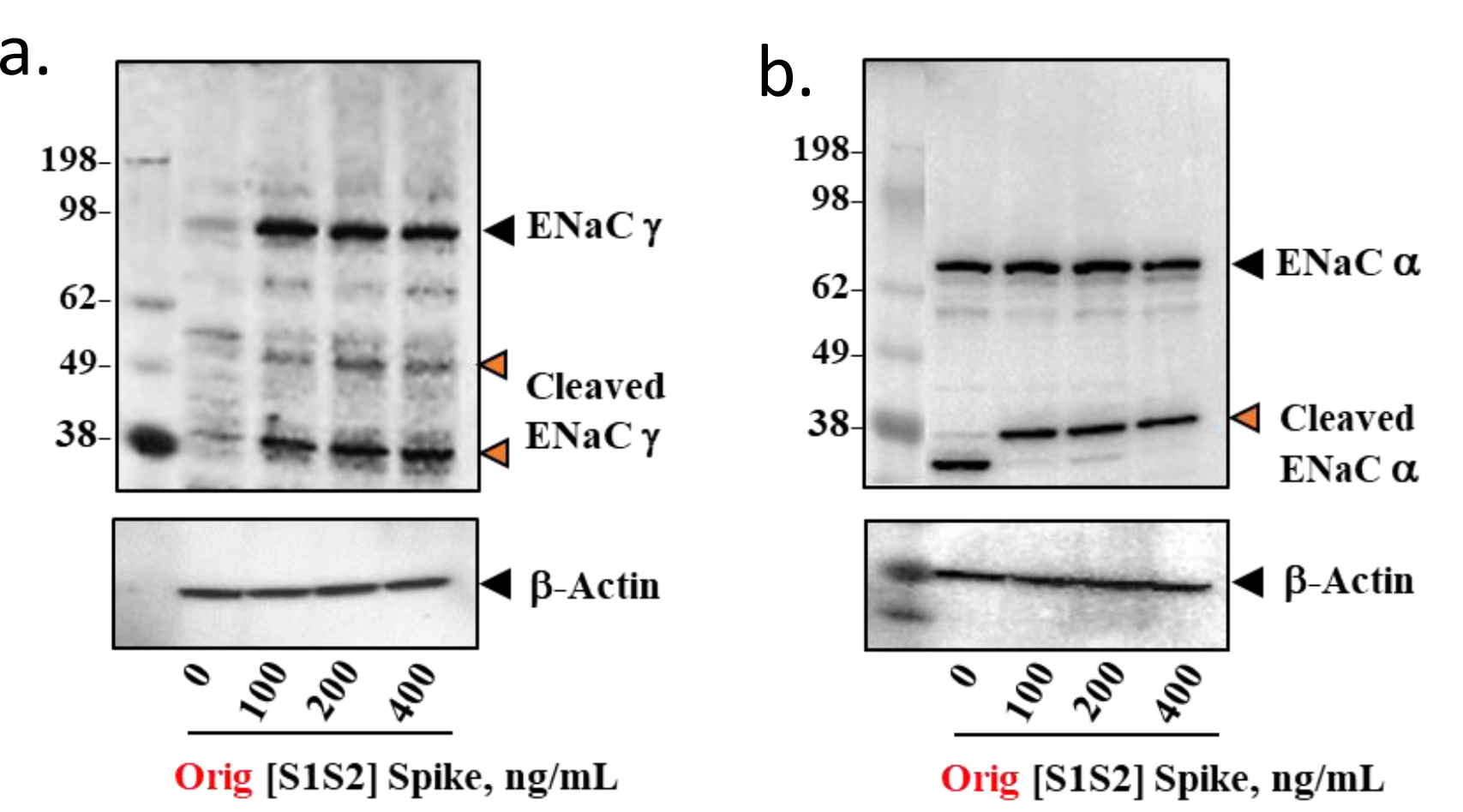
SARS-CoV-2 [S1S2] Spike Increases proteolytic activation of. α **and** γ **ENAC Subunits in Differentiated BCi-NS1.1 Cells. (a)** After Ussing chamber analyses, cells were washed with PBS and lysed in RIPA buffer supplemented with the anti-protease/phosphatase cocktail. Equivalent amounts of lysates (50 μg/sample) were electrophoresed on 4-12% or 4-20% gradient gels (Invitrogen), transferred to PVDF membranes, and membranes were probed with β-Actin antibody (Sigma) and an antibody against γ ENaC. **(b**) Epithelia were managed as described for **Part a**, and then probed with an antibody against α ENaC.

## Discussion

The mechanism for the proinflammatory phenotype of the COVID-19 airway has been an unsolved puzzle since the earliest days of the pandemic. Here we have used a model human small airway epithelium as an experimental platform, and from the data we are able to predict that inflammation in the COVID-19 airway may be due to SARS-CoV-2 Spike-dependent inhibition of CFTR protein expression. We find that ACE2 co-immunoprecipitates with CFTR, suggesting that ACE2 may be the link between Spike protein and CFTR. We also find that loss of CFTR protein can be traced to inability to recover plasma membrane-localized CFTR from the endocytic recycling process. Since the purpose of endocytic recycling is to eliminate damaged proteins, it is likely that the Spike protein, acting through ACE2, confers some form of damage or instability on the CFTR. Kinetic analysis suggests that the mechanism for Spike-dependent elimination of CFTR might depend on Spike first binding to ACE2. Consistently, the cardiac glycoside drugs ouabain, digitoxin and digoxin, which are competitive inhibitors of Spike protein binding to ACE2 ^20^, are able to rescue Spike-induced loss of cAMP-activated CFTR channel activity. In addition, from the perspective of inflammation, the loss of CFTR increased TNFα/NFκB signaling, as indicated by increases in TRADD. This is accompanied by increases in phosphorylated IκBα, and the chemokine IL-8. Finally, the loss of CFTR resulted in proteolytic activation of both α and γ ENaC. These affected pathways are depicted schematically in **Supplemental Figure S1.** *To our knowledge this is the first time COVID-19 airway inflammation has been experimentally traced to a contribution from SARS-CoV-2 Spike-dependent inhibition of CFTR signaling. This insight may have therapeutic implications since the cardiac glycosides, especially digitoxin and ouabain, are able to rescue CFTR channel activity from Spike-induced loss*.

### Clinical evidence for involvement of CFTR in COVID-19 susceptibility and outcome

In support of possible therapeutic implications of the findings summarized above, a reasonable question might be to what extent a reduction in CFTR expression in the lung, or elsewhere in CF-affected tissues, might relate in any way to COVID-19 disease in humans. One possibility would be to consider the parents of children with CF. Since CF is an autosomal recessive disease, these parents, also termed CF carriers, carry one mutant CFTR gene and one wildtype CFTR gene. The mutant CFTR gene does not contribute any functional CFTR protein. Unfortunately, the single wildtype gene in CF carriers does not compensate for the loss of the mutant CFTR gene. Consequently CF carriers suffer from reduction in CFTR function by approximately 50% ^45, 46^. Furthermore, recent large scale studies of CF carriers in Denmark ^47^ and the United States ^48^ show that CF carriers have a high risk of CF-related comorbidities, including respiratory tract infections, pancreatitis, hepatitis and others. CF carriers would thus appear to have a narrow functional window with respect to CFTR. It is therefore consistent with our results that in a recent study from the University of Siena, a cohort of hospitalized adult CF carriers with COVID-19 were reported to be more likely than non-CF carriers to develop a form of COVID-19 characterized by acute respiratory distress syndrome, high inflammatory response and early mortality by day 14 ^49^. The analysis included an adjustment for age, sex and comorbidities. Thus a reduction of CFTR in airway cells is a potent risk factor for adverse outcomes in COVID-19 disease. Consistently, a more recent geographical distribution study of COVID-19 spread and fatality showed that the incidence and severity of COVID-19 was proportionately increased in countries with the highest proportion of CF carriers in their population ^50^. *Based on our results, it is thus possible that exposure of airway cells in CF carriers to SARS-CoV-2 Spike protein may make residual wildtype CFTR protein even lower, thus contributing to an even worse outcome for CF carriers with COVID-19*.

### Epidemiological evidence for a therapeutic contribution to COVID-19 by cardiac glycoside drugs

Additional therapeutic implications are also implicit in the newly described ability of nanomolar concentrations of cardiac glycoside drugs to competitively inhibit Spike:ACE2 binding, and thus to block infectivity of lung cells by native SARS-CoV-2 ^20^. We recently approached this question by conducting a cross-sectional study of the 9.6 million patient U.S. Military Health System Data Repository (MDR), to identify all beneficiaries with a heart failure diagnosis during the COVID-19 pandemic period of April 1, 2020 to August 31, 2021 ^51^. We then identified all those who received standard-of-care, and those who because of increased comorbidities received digoxin instead. The latter patients were limited to digoxin because only digoxin is currently licensed for use in the United States. However, increased comorbidities, as defined by the Charlson Comorbidity Index (CMI) are widely described as being associated with worse outcomes for COVID-19 ^52–55^ . Nonetheless, digoxin-treated heart failure patients proved to be protected equivalently to those well enough to receive standard-of-care. *These data are thus consistent with the possibility that drugs in the class of cardiac glycosides might have a prospective therapeutic contribution to make if subjected to a clinical trial in the future*.

Although COVID-19 is a lethal disease, drug safety remains an issue for consideration. Fortunately, these drugs have been in human use since at least 1776, so much is known about their pharmacology ^56^. For example, there are upper concentration limits when treating cardiac patients with these drugs who suffer from heart failure or arrhythmias. Thus for patients in this category these drugs are said to have a narrow therapeutic index ^57^. However, for the vast majority of patients with normal hearts, ingestion “…of large but not lethal quantities of digitalis, either in an attempt at suicide or by accident, premature impulses and rapid arrhythmias are infrequent.” ^57^. Consistently, no drug-related adverse events have been reported in trials on subjects with normal hearts who were given clinical doses of ouabain ^58, 59^, digitoxin ^23, 60^, or digoxin ^61–64^. *Nonetheless, it is important that treatment with these drugs should always be under the control of health care professionals*.

### Renin-Angiotensin system and ENaC in the lung

Under normal conditions, ENaC activation in the lung is believed to be positively controlled by Angiotensin II, and negatively controlled by Angiotensin (1-7). Inversely, the presence of CFTR negatively controls ENaC in airway epithelial cells, while the absence of CFTR activates ENaC channels (see **Supplemental Figure S1**). ACE in the lung pulmonary vasculature converts Angiotensin I to Angiotensin II, the principal Renin-Angiotensin-Aldosterone System (RAAS) effector peptide, not only for the lung but for the entire circulation ^31^. Angiotensin II can then directly activate pulmonary ENaC, via the AT1 receptor, or inhibit pulmonary ENaC via the AT2 receptor ^32, 65^. However, angiotensin II is also inactivated by ACE2 on airway epithelial cells where the Angiotensin (1-7) product inhibits pulmonary ENaC via MSR and the AT2 receptor (see **Supplemental Figure S1**). There is thus a balance struck by ACE and ACE2 to regulate Angiotensin II concentration and therefore ENaC activity in the lung. On the other hand, CFTR appears to control passive ENaC activity by transporting negative chloride ions into the airway. To compensate for high levels of negative chloride ions in the airway, positive sodium ions enter the airway passively through the inactive ENaC channel. Together the Na^+^ and Cl^-^ ions passively draw water into the airway ^43, 66, 67^. ENaC is located mostly on ciliary cells in the lung, while CFTR is mostly expressed in secretory cells ^68^. Thus the need to compensate for excess negative charge in the airway may be the most likely basis for physiological control of the passive ENaC contribution to airway hydration on a cell-by-cell basis. The physiological consequence is a dynamic and responsive hydration process which helps to maintain the paraciliary fluid layer upon which mucins are propelled upward by cilia to either be expectorated or swallowed ^66^. However, in the absence of functional CFTR, either by mutation or by Spike-dependent loss, ENaC is activated by serine proteases, including TMPRSS2 and FURIN ^8, 69, 70^. *Excessive sodium absorption ensues: dehydrating the airway; disrupting mucociliary clearance; and activating inflammation* ^9, 18, 19, 43^.

It is at this point that the mechanisms of ENaC activation and the mechanism of viral invasion of airway epithelial cells by SARS-CoV-2 intersect ^8^. For the virus to enter the epithelial cell in the lung, the viral [S1S2] SARS-CoV-2 Spike protein must bind to ACE2. The Spike protein is then further proteolytically cleaved at the [S1S2] junction into the [S1 + S2] Spike protein by TMPRSS2, FURIN, and possibly other serine proteases ^71, 72^ (see **Figure 8** and **Supplemental Figure S1**). These proteolytic events are thus key to viral infectivity and development of COVID-19. Coincidentally these same proteases are evolutionarily located at the apical membrane surface for the physiological purpose of proteolytically activating the α and γ subunits of the heterotrimeric ENaC channel. It has been speculated that the SARS-CoV-2 virus has taken advantage of this existing protease system by evolving a CendR amino acid sequence at the S1S2 junction on the Spike protein that enables the existing TMPRSS2 and FURIN to drive infectivity ^8^. *The BCi-NS1.1 basal cells and differentiated epithelia both contain high levels of TMPRSS2 and FURIN* ^38^*, and the coincidence of these two processes in the epithelial culture is manifest by the Spike-dependent proteolytic activation of α and γ ENaC, as shown in* ***Figure 8***.

### Other pulmonary illnesses that involve loss of CFTR

Induced loss of CFTR by environmental or infectious agents has been observed in the past. For example, Chronic Obstructive Pulmonary Disease (COPD) is associated with smoke-induced, sustained loss of CFTR ^73–76^. Reduced CFTR mRNA levels are also seen in scrape biopsies from nasal epithelia of COPD patients ^73, 77^. The agent affecting CFTR in cigarette smoke has been identified as acrolein, which inhibits CFTR chloride channel gating, not only in the lung but also in the intestine ^78^. A second example is infection of HEK-293 cells by influenza strain A/Udorn, which has been reported to reduce CFTR chloride channel activity and CFTR protein ^79, 80^. The viral matrix protein M2 has been proposed to promote intracellular lysosomal degradation of CFTR, although the biochemical mechanism is not well understood. These examples thus suggest a similarity of disease states in which the CFTR protein is diminished by an external factor rather than a classical mutation. Thus *the reducing effect of SARS-CoV-2 Spike protein on CFTR* protein expression could be a third example.

### Limitations of the study

This study has limitations. First, it is a limitation of our study that we do not yet understand how ACE2 binds to CFTR. Nonetheless, we know experimentally that they do bind together, both in epithelia and in basal cells. Second, it is a limitation that we do not yet understand the nature of the apparent damage or instability inflicted on CFTR by the Spike:ACE2:CFTR interaction. Nonetheless, the classical impermeant biotin experiment indicates this must happen. Third, we do not yet know which cells in the model epithelia are experimentally responsible for the CFTR and ACE2 activities. However, the basal cells possess both ACE2 and CFTR and share some activities with the epithelia. Furthermore, secretory cells, which dominate airway CFTR expression in native superficial epithelia ^81^, are present in the differentiated BCi-NS1.1 epithelia ^39^, as is ACE2 ^38^. We suggest that substantively addressing these questions would be clearly beyond the experimental scope of the present study and look forward to addressing them in the future.

## Conclusion

Based on these investigations with a model of small airway epithelia we predict that increased TNFα/NFκB- and ENaC-dependent inflammation in the COVID-19 airway is mediated by inhibition of CFTR signaling by SARS-CoV-2 Spike protein, thus inducing a pro-inflammatory CFTR-null clinical phenotype in lung epithelia. Based on descriptions of more severe COVID-19 in adult CF carriers with only one copy of wildtype CFTR we suggest that this model-based conclusion may be consistent with patient-based experience.

## Methods

### Cells and culture conditions

As a cellular platform for the study we have used the hTERT-transformed BCi-NS1.1 basal stem cell differentiated on the air-liquid-interface (ALI) ^38, 39^. We thank Dr. R.G. Crystal (Cornell Medical College, New York City, NY) for the gift of the cells. The cells used here came as Passage 11 following original derivation. The serum-free culture medium was used directly as indicated (Walters et al, 2013). Differentiation in the ALI was conducted over a 25 day period exactly as described ^39^. Cyclic AMP-activated CFTR chloride channel activity was measured using a Ussing Chamber as previously described ^23^.

### Co-immunoprecipitation of ACE2 and CFTR

At the end of an experiment media were removed; epithelia or cells were washed with PBS; and then lysed in RIPA buffer containing protease and phosphatase inhibitors. Total lysates were precleared with Protein A Dynabeads (Thermo Scientific), incubated for 24 hr at 4°C with 2 μg of normal rabbit serum (NRS) or rabbit anti-ACE2 antibody (proteintech), and finally incubated for 4 hr with Protein A Dynabeads. CFTR in the immuno-precipitated protein complexes was analyzed by western blotting using anti-CFTR antibodies, clones UNC 570 and UNC 660, development of which were which were funded by the Cystic Fibrosis Foundation, and purchased by us from the University of North Carolina, Chapel Hill, NC). See ^21^ for further details.

Electrophysiology of epithelial cultures at the air-liquid-interface (ALI).

Human hTERT-transformed BCi-NS1.1 basal stem cells were seeded at a concentration of 4.5 x 10^5^ cells/cm^2^ on Transwell inserts (0.4 mm size pore; Corning Inc, NY) and differentiated at the air-water interface for 25 days ^39^. Cells were exposed to different concentrations of Spike protein for 4 hours, washed with fresh medium, and then incubated for a further 24 hours. Snapwell inserts were mounted in Ussing Chambers (Physiologic Instruments, Reno Nevada). A basolateral-to-apical chloride gradient was imposed by partially replacing NaCl with NaGluconate. The epithelium was voltage clamped and short-circuit current (Isc) was measured. CFTR channel activity was detected by first inactivating sodium currents with 100 μM amiloride; then activating CFTR chloride channels with 10 μM forskolin and 100 μM IBMX; then specifically inactivating CFTR channels with 10 μM CFTRinh-172. See ^23, 82^ for further details.

### Measurement of ACE2 and CFTR endosomal recycling

Apical surfaces of epithelia were treated with different concentrations of Spike protein for 4 hours; and then cells were incubated for a further 20 hours at 37°C in ALI conditions. Cells were then washed with ice cold Ca^+2^/Mg^+2^ PBS buffer 3 times and then incubated for an additional 24 hr at 4°C with 2mg/ml impermeant biotinylation reagent, sulfo-NHS-S-S biotin (Thermo Scientific). Biotin-labeled cells were lysed in RIPA buffer, and biotinylated proteins were extracted using Streptavidin beads (Thermo Scientific). Levels of cell surface CFTR in the bead extract were finally analyzed by western blotting. CFTR antibodies (Clone 570 and 660) were purchased from the North Carolina Cystic Fibrosis Center (funded by the Cystic Fibrosis Foundation, Bethesda, MD).

### Measurement of IL-8

Well differentiated BCi-NS1.1 cells on 12mm Snapwell inserts were treated for 4 hours apically with media or various concentrations of Original-[S1S2] Spike protein. The conditioned media in the apical side were then removed. Cells were washed with culture media and continued to incubate further for 20 hr under ALI conditions. The concentration of IL-8 released in basolateral culture media were assayed by ELISA using the human ELISA development kits (DuoSet DY208, R&D Systems, Minneapolis, MN, USA) according to the manufacturer’s instructions.

### Chemicals and biologics

Digitoxin, digoxin, and ouabain were purchased from Sigma-Aldrich. Drugs and derivatives were solubilized as 4 mM stock solutions in 100% DMSO, and aliquots were serially diluted in the same solvent, and then into assay medium. In assays containing DMSO, the final solvent concentrations were 0.01% or less. With respect to identifying recombinant Spike protein sequences, we have adopted the World Health Organization (WHO) naming system as follows:

(i) The Spike protein from the original SARS-CoV-2 virus, which expressed aspartic acid at position 614 (*viz*., [D614]) was labeled as Orig-[S1S2] Spike or Original-[S1S2] Spike. (ii) The mutant β-Spike 1.315 was labelled as β-1.315 [S1S2] Spike. Recombinant proteins produced in human T293 cells were obtained as follows: Recombinant (Original, Wuhan) SARS-CoV-2 Spike [S1+S2] protein was expressed in Baculovirus insect cells with polyhistidine tag at the C-terminal (Cat # 40589-V08B1), and β-1.315 [S1+S2] protein (K417N, E484K, N501Y, D614G, A701V-His; Cat # 40589-V08811) were obtained from SinoBiologicals (Philadelphia, PA).

### Statistics

To determine the significance of changes in kinetic parameters of ACE2 binding to SARS-CoV-2 Spike mutants we applied least-squares regression to the linearized Eadie-Hoffstee plot of the binding data, and determined the statistical significance of the difference between the slopes (*i.e*., KD’s) and between the intercepts (*i.e*., Bmax) using R or Stata statistical packages. Ki values were calculated for each data point in the linear range depending on the inhibition mechanism. Except as noted, all plotted data points are the means of 3 or more independent experiments. Means ± SE were calculated; p <0.05 were taken to indicate a significant difference from controls.

## Acknowledgements

We gratefully acknowledge support for this project by the Cooperative Health Initiative Research Program (CHIRP), supported by NHLBI/NIH (IAA-AA-HL-14-007; H.B. Pollard, PI); and by the Consortium for Health and Military Performance (CHAMP), supported by Warfighter Readiness: Optimizing Human Performance (HU00011920047; MEM-91-10314; P. Deuster, PI); by the Center for the Study of Traumatic Stress (CSTS), supported by DOD Core Grant (HU00012120068, R. Ursano, PI.), and by Intramural Research Program Award Grant No. APG-70-12301 (H.B. Pollard, PI). We thank Laiman Tavedi for expert technical and administrative support. We thank Alan C-Y Hsu for collegial helpful discussions. The opinions, interpretations, conclusions and recommendations are those of the authors and are not necessarily endorsed by the U.S. Army, Department of Defense, the U.S. Government or the Uniformed Services University of the Health Sciences. The use of trade names does not constitute an official endorsement or approval of the use of such reagents or commercial hardware or software. This document may not be cited for purposes of advertisement.

## Contributions

HC, HP, OE, QY, and BP designed experiments. HC, QY, and TC performed experiments. OE, NW, HC, TC and HP contributed to data analysis. HP, HC, OE, TC and BP and NW, wrote the paper. All co-authors critically read the paper.

## Supplemental Data

### Supplemental Methods

#### Enzyme Linked Immunosorbant Assay (ELISA) for interaction between Spike proteins and ACE2

Purified recombinant SARS-CoV-2 Spike proteins were individually dissolved in coating buffer (16 mM Na2CO3, 34 mM NaHCO3, pH 9.6) at a concentration of 2 μg/ml Spike protein. This solution, in 100 μL aliquots, was then added to wells of Costar 96 well plates (Corning, Corning, NY; Catalog # 2592) and incubated overnight at 4°C. The next day, wells were washed 3 times in Phosphate Buffered Saline with 0.05% Tween 20 (PBST) at room temperature. Wells were then blocked with 300 μL Blocking Buffer (0.5% Bovine Serum Albumin (BSA), dissolved in PBST) for two hours at 37°C. Wells were then washed 3 times in PBST at room temperature (68°F). When cardiac glycosides were to be added, they were dissolved in Reagent Diluent (0.5% BSA in PBST), added in 100 μL volumes to each well, and incubated at 37°C overnight. Further steps were as described in our recent publication ^20^. The final results are given as averages ± Standard Errors of all independent experiments.

#### Recombinant proteins and antibodies

Recombinant proteins produced in human T293 cells were obtained as follows: Recombinant (exodomain) Human Angiotensin Converting Enzyme 2 (ACE2) (Cat # 230-30165; lot 04U0620TW) was purchased from RayBiotech (Peachtree Cornewrs, GA, 30092). Human ACE2 Biotinylated Antibody (Cat # BAF933) was obtained from R&D Systems (Minneapolis, MN, 55413).

### Supplemental Figures

**Figure S1.**
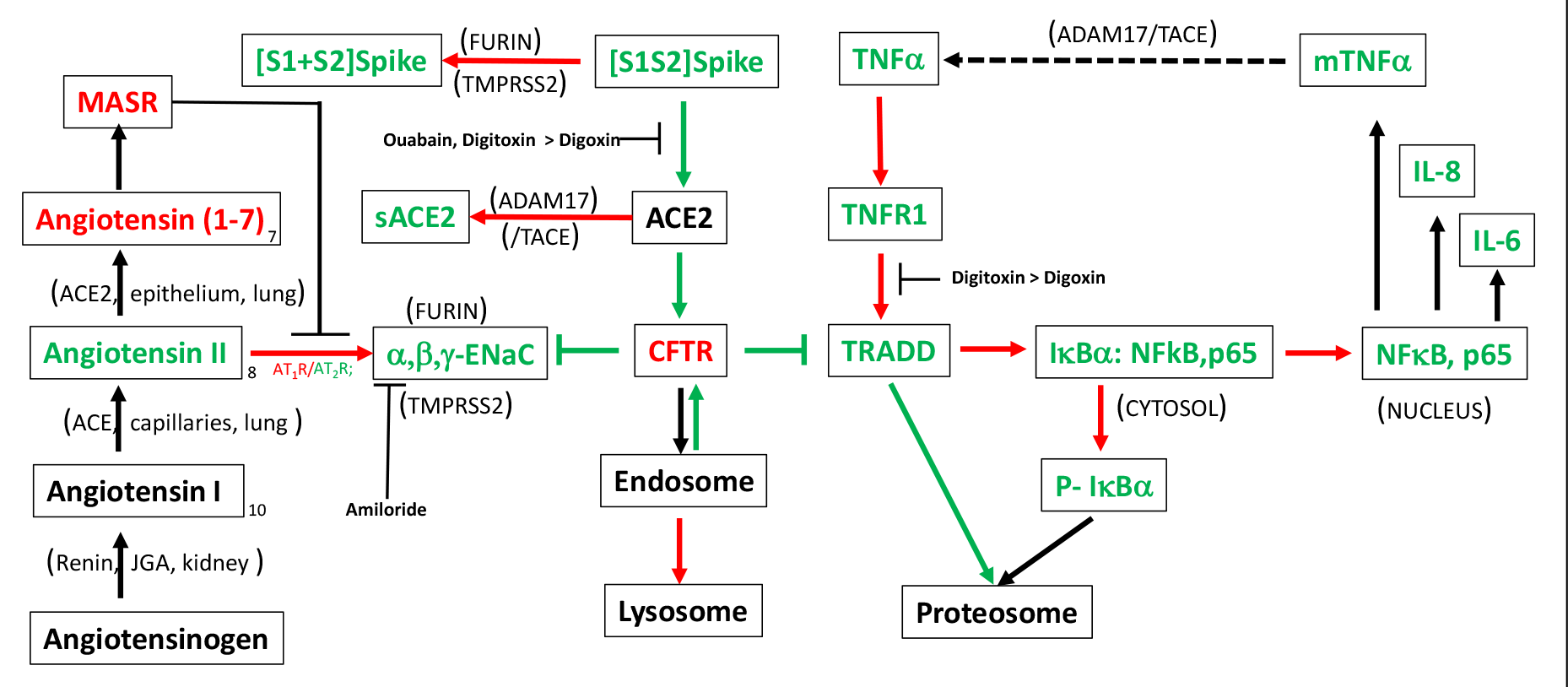
**Inactivation of CFTR by Spike protein activates both NFkB,p65 and α,b,g ENaC signaling**. Under baseline conditions CFTR tonically suppresses ENaC and TRADD. However, upon addition of [S1S2] Spike, CFTR is dose-dependently reduced by failure to be recovered from endosomal recycling (this paper). In the absence of CFTR, TRADD is no longer constitutively directed to the proteosome, and it is free to activate IKKαβγ, which phosphorylates IκBα. NFκB, p65 is now free to leave the cytosol and enter the nucleus. Cytokines and chemokines such as IL-6, IL-8 and mTNFα are then expressed. Membrane-bound TNFα (mTNFα) is converted to soluble sTNFα/TNFα by ADAM17/TACE). ADAM17/TACE also converts membrane bound mACE2 to soluble sACE2. Cardiac glycoside drugs ouabain, digitoxin and digoxin are potent competitive inhibitors of Spike:ACE2 binding (Caohuy H. et al, Scientific Reports,2021). Digitoxin separately blocks interactions in the host between the TNFα/TNFR1 complex and TRADD (Yang Q et al, PNAS, 2005). In the absence of CFTR, ENaC is also proteolytically activated by FURIN and TMPRSS2 (this paper). In preparation for penetration into the target cell, the same proteases cleave Spike at the S1S2 junction. Color code: **red** = elevated; **green** = reduced; black = no known change.

**Figure S2.**
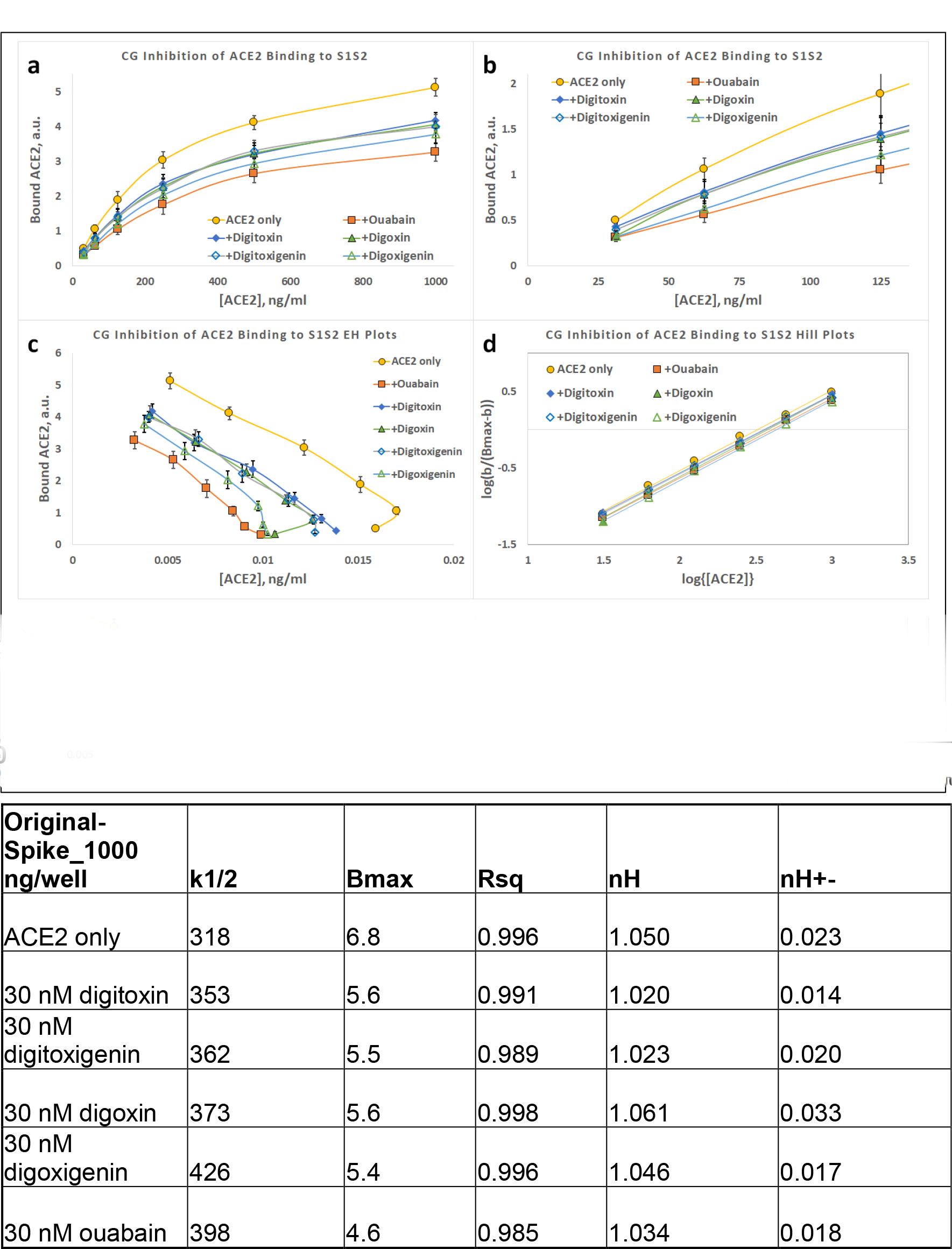
Binding of ACE2 to Original-Spike protein. (**a**) Substrate-Binding plots for ACE2 binding to Original [D614] α-Spike in presence and absence of cardiac glycoside drugs. **(b)** Substrate-Binding plot for lower concentrations of ACE2. Note concave-down structure at low binding levels. (**c**) Eadie-Hoffstee plots of data in Part (**a**). (**d**) Hill plots for data in Part (**a**). Each point is the average ± SE for N= 5-6 independent experiments.

**Supplemental Figure S3.**
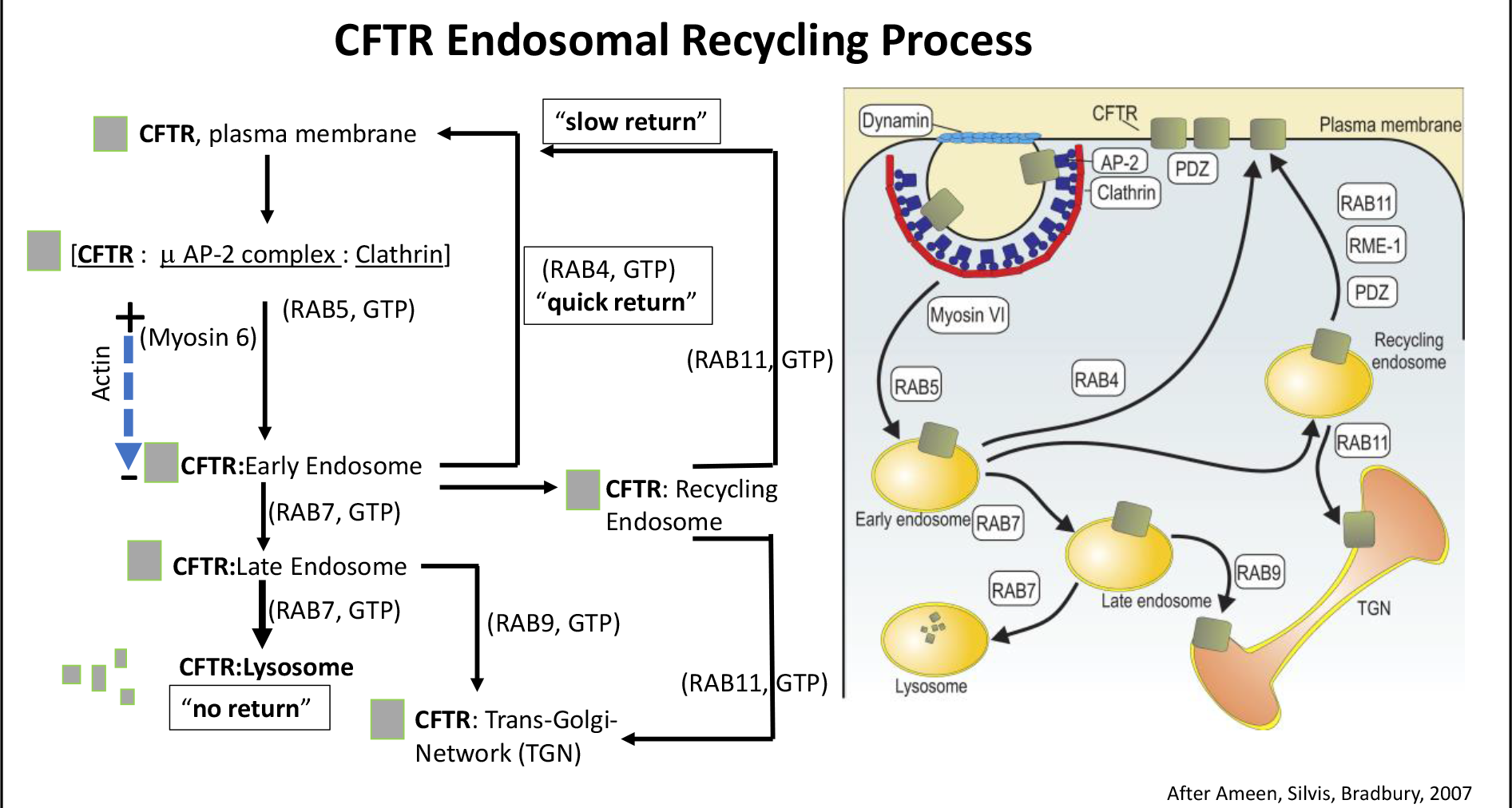
CFTR Endosomal Recycling Process. CFTR proteins on the epithelial cell surface are internalized by a Clathrin/ Dynamin-dependent process that depends on the Ras-related protein RAB5 for movement of CFTR to early endosomes. Myosin 6 drives physical internalization by binding to the (+) end of actin and moving cargo towards the (-) actin end. RAB5 recruits RAB7 to drive maturation of early endosomes to late endosomes. Maturation includes transporting vacuolar (H+) ATPases (V-ATPases) from the Trans-Golgi Network (TGN) to the endocytic vesicles. RAB4 mediates quick return of CFTR-laden endosomes to the plasma membrane. RAB11 mediates slower return to the plasma membrane. Late endosomes are also conveyed to the lysosome for destruction of “damaged” CFTR. SARS-CoV-2 Spike protein drives destruction of cell surface CFTR by this endosomal recycling process. RAB9 mediates transfer of CFTR from Late endosome to the Trans-Golgi-Network (TGN). Grey squares represents CFTR. Fragmented grey squares represents proteolysis of CFTR in the lysosome. Cartoon on the right is from Ameen, Silvis and Bradbury, 2007. Pathways on the left represent assay-based identified transitions.

